# PaleoProPhyler: a reproducible pipeline for phylogenetic inference using ancient proteins

**DOI:** 10.1101/2022.12.12.519721

**Authors:** Ioannis Patramanis, Jazmín Ramos-Madrigal, Enrico Cappellini, Fernando Racimo

## Abstract

Ancient proteins from fossilized or semi-fossilized remains can yield phylogenetic information at broad temporal horizons, in some cases even millions of years into the past. In recent years, peptides extracted from archaic hominins and long-extinct mega-fauna have enabled unprecedented insights into their evolutionary history. In contrast to the field of ancient DNA - where several computational methods exist to process and analyze sequencing data - few tools exist for handling ancient protein sequence data. Instead, most studies rely on loosely combined custom scripts, which makes it difficult to reproduce results or share methodologies across research groups. Here, we present PaleoProPhyler: a new fully reproducible pipeline for aligning ancient peptide data and subsequently performing phylogenetic analyses. The pipeline can not only process various forms of proteomic data, but also easily harness genetic data in different formats (CRAM, BAM, VCF) and translate it, allowing the user to create reference panels for phyloproteomic analyses. We describe the various steps of the pipeline and its many functionalities, and provide some examples of how to use it. PaleoProPhyler allows researchers with little bioinformatics experience to efficiently analyze palaeoproteomic sequences, so as to derive insights from this valuable source of evolutionary data.

## Introduction

Recent advances in protein extraction and mass spectrometry [1, 2, 3, 4] have made it possible to isolate ancient peptides from organisms that lived thousands or even millions of years ago. Certain ancient proteins have a lower degradation rate and can be preserved for longer than ancient DNA [5, 6, 7, 8]. These ancient proteins can be utilized by Peptide Mass Fingerprinting (PMF) methods [9], including ZooMS [10], for genus or species identification [11] and to single out fossil material of interest for further analyses including DNA sequencing [12, 13], radiocarbon dating [14] and shotgun proteomics [12, 15]. Shotgun proteomics in particular, utilizing tandem mass spectrometry, has enabled the reconstruction of the amino acid sequences of those proteins, which sometimes number in the hundreds [16, 17]. These sequences contain evolutionary information and thus have the potential to resolve important scientific questions about the deep past, which are not approachable via other methods. Tooth enamel proteins and bone collagen in particular have been successfully extracted from multiple extinct species, in order to resolve their relationships to other species [18, 19, 20, 21, 22, 23, 24, 25, 26].

Ancient proteomic studies typically use combinations of custom scripts and repurposed software, which require extensive in-house knowledge and phylogenetic expertise, and are not easily reproducible. Barriers to newcomers in the field include difficulties in properly aligning the fractured peptides with present-day sequences, translating available genomic data for comparison, and porting proteomic data into standard phylogenetic packages. The creation of automated pipelines like PALEOMIX [27] and EAGER [28] have facilitated the streamlining and reproducibility of ancient DNA analyses, which has been particularly helpful for emerging research groups around the world. This has undoubtedly contributed to the growth of the field [29]. Yet, the field of palaeoproteomics still lacks a “democratizing” tool that is approachable to researchers of different backgrounds and expertises.

Another important issue in phyloproteomics is the relative scarcity of proteomic datasets [30, 31]. There are currently tens of thousands of publicly available whole genome sequences, covering hundreds of species [32, 33, 34, 35, 36]. The amount of publicly available proteome sequences is much smaller. NCBI’s list of sequenced genomes [37] includes 78,420 species, out of which 30,530 are eukaryotes and 11,345 labeled as ‘Animal’. For com-parison, Uniprot’s reference proteomes list [38] contains a total of 23,805 entries of which 2,400 are eukaryotes and around 950 are labeled as ‘Metazoa’. For most vertebrate species, lab-generated protein data does not exist and phyloproteomic research is reliant on sequences translated *in silico* from genomic data. These, more often than not, are not sufficiently validated or curated [39]. Ensembl’s curated database of fully annotated genomes, and thus available proteomes, numbers only around 270 species [40]. As a result, assembling a proper reference dataset for phyloproteomics can be challenging. Given how important rigorous taxon sampling is in performing proper phylogenetic reconstruction [41, 42], having a complete and reliable reference dataset is crucial. In the case of proteins, the typically short sequence length and the low amounts of sequence diversity - due to the strong influence of purifying selection - means that absence of knowledge about a single amino acid polymorphism (SAP) can strongly affect downstream inferences [23, 43, 44, 45].

## Methods

To address all of the above issues, we present “Paleo-ProPhyler”: a fully reproducible and easily deployable pipeline for assisting researchers in phyloproteomic analyses of ancient peptides. “PaleoProPhyler” is based on the workflows developed in earlier ancient protein studies [22, 24, 26], with some additional functionalities. It allows for the search and access of available reference proteomes, bulk translation of CRAM, BAM or VCF files into amino acid sequences in FASTA format, and various forms of phylogenetic tree reconstruction.

To maximize reproducibility, accessibility and scalability, we have built our pipeline using Snakemake [46] and Conda [47]. The Snakemake format provides the workflow with tools for automation and computational optimization, while Conda enables the pipeline to operate on different platforms, granting it ease of access and portability. The pipeline is divided into three distinct but interacting modules (Modules 1,2 and 3), each of which is composed of a Snakemake script and a Conda environment 1.

Module 1 is designed to provide the user with a baseline (curated) reference dataset as well as the resources required to perform the in silico translation of proteins from mapped whole genomes. The input of module 1 is a user-provided list of proteins and a list of organisms. The user also has the option of choosing a particular reference build. Utilizing the Ensembl API [48], the module will return 3 different resources for each requested protein and for each requested organism. These are : a) the reference protein sequence of that organism in FASTA format [49], b) the location (position and strand) of the gene that corresponds to that protein and c) the start and end of each exon and intron of that gene / isoform. The downloaded FASTA sequences are available individually but are also assembled into species- and protein-specific datasets. They can be immediately used as a reference dataset for either downstream phylogenetic analyses or as an input database for mass spectrometry software, like MaxQuant [50], Pfind [51], PEAKS [52] and others [53, 54, 55, 56]. The gene location information and the exon / intron tables can be utilized automatically by Module 2. For each requested protein, the module will select the Ensembl canonical isoform by default. Should the user desire a specific isoform or all protein coding isoforms of a protein, they have the ability to specify that as an option in the provided protein list.

**Figure 1:**
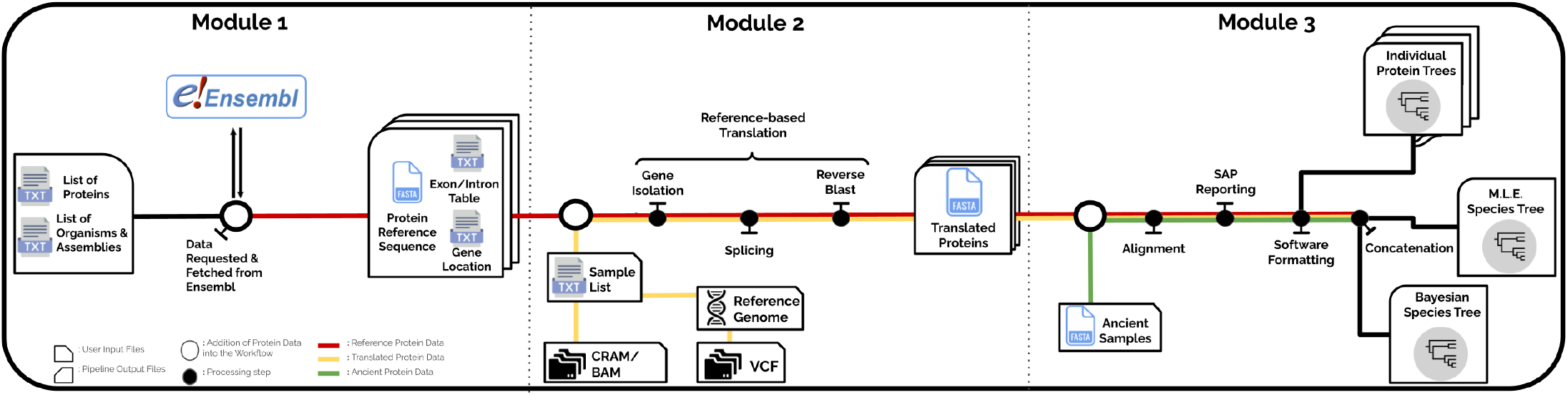
Overview of the Pipeline

Module 2 is designed to utilize the resources generated by Module 1 and to extract, splice and translate genes from whole genome data, into the proteins of interest. This module can handle some of the most commonly used genomic data file formats, including the BAM [57], CRAM [58] and VCF [59] formats. The easiest way to run Module 2 is to first run Module 1 for a set of proteins and a selected organism. This will generate all the necessary files and resources required for the protein translation. The selected organism will be used as a reference for the translation process. All genomic data to be translated must be mapped onto the same reference organism. The user can then run Module 2 simply by providing the organism’s name (and optionally a reference version), as well as a list of the samples to be translated. The user can also translate samples from a VCF file, but they will need to provide a reference genome in FASTA format, to complement the variation-only information of the VCF file. The translated protein sequences are available individually but are also assembled into individual- and protein-specific datasets.

Module 3 is designed to perform a phylogenetic analysis, with some modifications needed when working with palaeo-proteomic data. The input of this module is a FASTA file, containing all of the protein sequences from both the reference dataset and the ancient sample(s) to be analyzed. The dataset is automatically split into protein specific sub-datasets, each of which will be aligned and checked for Single Amino acid Polymorphisms (SAPs). The alignment is a two step process which includes first isolating and aligning the modern/reference dataset and then aligning the ancient samples onto the modern ones using Mafft [60]. Isobaric amino acids that cannot be distinguished from each other by some mass spectrometers are corrected to ensure the downstream phylogenetic analysis can proceed without issues. Specifically, any time an Isoleucine (I) or a Leucine (L) is identified in the alignment, all of the modern sequences are checked for that position. If all of them share one of the 2 amino acids, then the ancient samples are also switched to that amino acid. If both I and L appear on some present-day samples, both present-day and ancient samples are switched to an L. The user also has the option to provide an additional file named ‘MASKED’. Using this optional file, the user can mask a present-day sample such that it has the same missing sites as an ancient sample. Finally a small report is generated for each ancient sample in the dataset, and a maximum likelihood phylogenetic tree is generated for each protein sub-dataset through PhyML [61]. All protein alignments are then also merged together into a concatenated dataset. The concatenated dataset is used to generate a maximum-likelihood species tree [62] through PhyML and a Bayesian species tree [63, 64] through MrBayes [65] or RevBayes [66]. The tree generation is parallelized using Mpirun [67].

The modules are intended to work with each other, but can also be used independently. An in depth explanation of each step of each module, as well as the code being run in the background, is provided on the software’s Github page as well as in the supplementary material.

## Applications

As proof of principle, we deploy this pipeline in the reconstruction of ancient hominid history using the publicly available enamel proteomes of *Homo antecessor* and *Gigantopithecus blacki*, in combination with translated genomes from hundreds of present-day and ancient hominid samples. In the process, we have generated the most complete and up to date, molecular hominid phyloproteomic tree 2. The process of generating the reference dataset and its phyloproteomic tree using PaleoProPhyler is covered in detail in the step-by-step [Github Tutorial](https://github.com/johnpatramanis/Proteomic_Pipeline/blob/main/GitHub_Tutorial/Tutorial.md). The dataset used as input for the creation of the phylogenetic tree is available at [Zenodo](https://zenodo.org/record/7404802#.Y48vOHbMKbg)

## Protein Reference Dataset

In order to facilitate future analyses of ancient protein data, we also generated a publicly-available palaeoproteomic hominid reference dataset, using Modules 1 and 2. We translated 176 publicly available whole genomes from all 4 extant Hominid genera [33, 34, 68]. Details on the preparation of the translated samples can be found in the supplementary materials. We also translated multiple ancient genomes from VCF files, including those of several Neanderthals and one Denisovan [69, 70]. Since the dataset is tailored for palaeoproteomic tree sequence reconstruction, we chose to translate proteins that have previously been reported as present in either teeth or bone tissue. We compiled a list of 1.696 proteins from previous studies [71, 72, 73, 74, 75, 76] and successfully translated 1.543 of them. For each protein, we translated the canonical isoform as well as all alternative isoforms, leading to a total of 10.058 protein sequences for each individual in the dataset. Details on the creation of the protein list can be found in the supplementary materials. The palaeoproteomic hominid reference dataset is publicly available online at Zenodo, under the name ‘Hominid Palaeoproteomic Reference Dataset’

**Figure 2:**
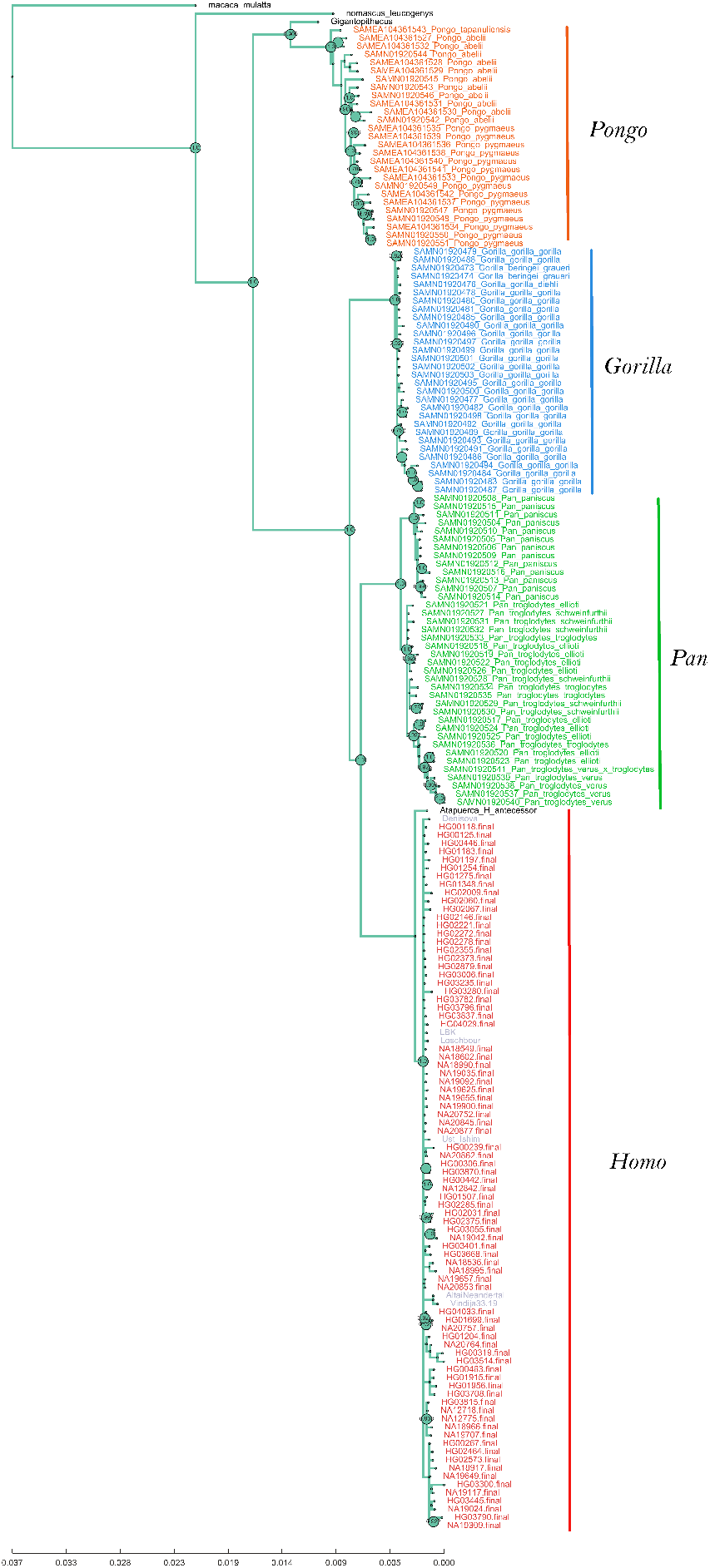
Phyloproteomic tree generated using PaleoProPhyler’s Module 3. The tree was constructed using 9 protein sequences obtained from enamel and includes more than 100 hominid individuals translated from genomic data, two individuals from published palaeoproteomic datasets as well as sequence data from a *Macaca* and a *Hylobates* individual, which are used to root the tree.

## Closing remarks

The workflows presented here aim to facilitate phylogenetic reconstruction using ancient protein data to a wider audience, as well as to streamline these processes and enable greater reproducibility in the field. Although we highly encourage the use of the tools and methods utilized by our workflows, we still caution against the over interpretation of palaeoproteomic results. Deriving species relationships from ancient proteins is still a relatively new endeavor and as a result, our understanding of this data, their quantity and quality requirements, robustness and accuracy are all largely unexplored. We believe that palaeoproteomic data should therefore be used in combination with other sources of information in order to make accurate evolutionary inferences.

## Author Contributions

- **Ioannis Patramanis**: Conceptualization, manuscript writing, code writing for the Snakemake scripts, compilation of the Conda environments and application of the pipelines to produce the results described in the ‘Application’ and ‘Protein Reference Dataset’ section.
- **Jazmin Ramos Madrigal**: Manuscript review, conceptualization and code for multiple R and bash scripts utilised by the Snakemake script as steps of the pipeline.
- **Enrico Cappellini** : Manuscript review and edit-ing
- **Fernando Racimo** : Conceptualization, manuscript writing, review and editing

## Acknowledgements

We thank Ryan Sinclair Paterson, Graham Gower, Alberto Taurozzi, Martin Petr, Evan Irving-Pease and other members of the Racimo and Cappellini groups, who provided valuable help and feedback throughout the project. We also thank Helen Fewlass for testing and identifying errors in the workflow.

## Data Availability

The Protein Reference Dataset is available on Zenodo: https://zenodo.org/record/7728060

PaleoProPhyler is publicly available on Github: https://github.com/johnpatramanis/Proteomic_Pipeline

The tool requires a Linux OS (Operating System) and the installation of Conda. The github repository contains a tutorial for using the workflow presented here, with the proteins recovered from the *Homo antecessor* and *Gigantopithecus blacki* as examples. We welcome code contributions, feature requests, and bug reports via Github. The software is released under a CC-BY license.

## Funding

The project was funded by the European Union’s EU Framework Programme for Research and Innovation Horizon 2020, under Grant Agreement No. 861389 - PUSHH. FR was additionally supported by a Villum Young Investigator Grant (project no. 00025300), a COREX ERC Synergy grant (ID 951385) and a Novo Nordisk Fonden Data Science Ascending Investigator Award (NNF22OC0076816). E.C. was additionally supported by the European Research Council (ERC) through the ERC Advanced Grant “BACKWARD”, under the European Union’s Horizon 2020 research and innovation program (grant agreement No. 101021361).

references

