## Supplementary Information for "PaleoProPhyler: a reproducible pipeline for phylogenetic inference using ancient proteins"

### PaleoProPhyler, Supplementary Material

### Detailed Overview of Pipelines

In the next few pages you will find a more in depth, step by step, overview of each module of the PaleoProPhyler workflow.

### Module 1

##### Description of Module:

**Module 1** is designed to provide the user with a baseline (curated) reference dataset as well as the resources required to perform the *in silico* translation of proteins from mapped whole genomes. The input of this module is a user-provided list of proteins and a list of organisms. Both input lists should be in a simple *.TXT file. The user also has the option of choosing a particular reference build. Utilizing the Ensembl API [1], the module will return 3 different resources for each requested protein and for each requested organism / reference build. The 3 output resources are:

1. The reference protein sequence in FASTA format [2].
2. The location (position and strand) of the gene that corresponds to the protein.
3. The start and end of each exon and intron of that gene/isoform.

The downloaded FASTA sequences are available individually but can also be assembled into species- and protein-specific datasets. They can be immediately used as a reference dataset for either downstream phylogenetic analyses or as an input database for mass spectrometry software, like MaxQuant [3], Pfind [4], PEAKS [5] and others [6, 7, 8, 9]. The gene location information and the exon/intron tables can be utilized automatically by Module 2. For the requested proteins, the module will select the Ensembl canonical isoform by default. Should the user desire a specific isoform or all protein coding isoforms of a protein, they have the ability to specify that as an option in the protein list *.TXT file.

##### Step by step scripts that run:

1. Python3 Script to fetch Ensembl Gene ID for each Protein - Organism Combination (Link to Script)
2. Python3 Script to fetch Transcript ID ( Ensembl-Canonical and Optional Alternative Isoforms ) for each Gene ID (Link to Script)
3. Python3 Script to fetch Protein Fasta Sequence for each Transcript ID (Link to Script)
4. Python3 Script to fetch Protein Exon and Intron information for each Transcript ID ( and create a table for them) (Link to Script)
5. Python3 Script to fetch Gene location (corrected for Reference version) for each Transcript ID (Link to Script)
6. Python3 Script to combine all FASTA Sequences into datasets ( per Protein and per Organism ) (Part of main Snakemake Script)

#### Input of Module 1:

- A TXT file with a list of coded protein names, one per line e.g.:

  AMELX
  ENAM
  AMELY

- Alternatively this list can also contain the names of specific isoforms, or the word ALL to bring all known isoforms e.g.:

  AMELX::AMELX-201
  ENAM::ENAM-202
  AMELY::ALL

  Example file here

- A TXT file with a list of scientific organism names e.g.:

  homo sapiens
  pan troglodytes
  gorilla gorilla

- Alternatively this second list can also contain the names of specific reference versions e.g.:

  homo sapiens GRCh37
  pan troglodytes CHIMP2.1.4
  gorilla gorilla gorGor3.1

Example file here

### Module 2

#### Description of Module:

**Module 2** is designed to utilize the resources generated by Module 1 and to extract, splice and translate genes from whole genome data, into the proteins of interest. Module 2 can handle some of the most commonly used genomic data file formats, including the BAM [10], CRAM [11] and VCF [12] formats. The easiest way to run Module 2 is to first run Module 1 for a set of proteins and a selected organism. This will generate all the necessary files and resources required for protein translation. The selected organism will be used as a reference for the translation process. All genomic data to be translated must be mapped onto the same reference organism. The user can then run Module 2 simply by providing the organism's name (and reference version), as well as a list of the samples to be translated, both in a *.TXT file. Should the user want to translate samples from a VCF file, they will also need to provide a reference genome in FASTA format, to complement the variation information of the VCF file. After executing the module, an initial 'normalization' step is performed, where all input files are formatted and indexed. Once this is complete, the locations of the genes are used to extract their sequence and the exon/intron information is used to splice them. These isolated and spliced genomic sequences are then BLASTed [13] onto the reference protein sequence, and the matching translated amino acids are stitched together into the final translated protein sequences. In the last step, the translated sequences are organized into 3 alternative databases:

1. The 'Per protein' database: a folder containing one FASTA file for each translated protein. Each protein FASTA file contains the sequences of that protein for all samples
2. The 'Per individual' database: a folder containing one FASTA file for each translated sample / individual. Each sample FASTA file contains all of the translated proteins for that sample.
3. The 'All Protein Reference' database: a single FASTA file containing all translated proteins for all samples.

Any of these FASTA files can be instantly merged with an ancient protein dataset and used in Module 3.

#### Input of the module:

This module has 3 alternative input file options:

- BAM input:
  - Samples to be translated should be named inside a .TXT file named 'Samples.txt'. The name of each sample should correspond to the name of the BAM file , but without the ending (e.g. '.bam') Example File. The BAM files themselves should be placed within your

    `"Dataset_Construction/Workspace/1_OG_BAM_FILES/"` folder.
  - Headers are renamed, e.g. from 'Chr1' to just '1', using the commands `'samtools view'` and `'samtools reheader'`
  - Renamed BAM file is index with `'samtools index -b'`
  - BAM file is split into chromosome BAM files using `'samtools view -b'`
  - Chromosome BAM files are re-indexed using `'samtools index -b'`
  - BAM files are transformed to FASTA file using

"angsd -minQ 30 -minMapQ 30 -doFasta 2 -doCounts 1 . For heterozygous positions only one of the two alleles is represented, with the one with the highest number of reads being chosen and a random when the counts are tied. -basesPerLine 60"

- CRAM input:
  - Input should have the same format as described in the BAM files
  - Cram files are transformed internally into BAM file using the command `"samtools view -b"`
  - Follows exactly the same input path as the BAM files above.
- VCF input:
  - Samples to be translated should be named inside a .TXT file named

    `'VCF_Samples.txt'`. The name of each sample should be written as it exists within the VCF file. In addition the name of the VCF file needs to be provided , as well as the name of the reference Fasta file (see below) Example File. The VCF files themselves should be placed within your

    `'Dataset_Construction/Workspace/0_VCF_FILES/'` folder.
  - A reference genome in FASTA format needs to be provided and placed in the

    `'Dataset_Construction/Reference/'` folder.
  - The provided reference genome is renamed to the same standard as the BAM files ('Chr1' to just '1')
  - The VCF file is renamed as well, using `'bcftools view'` , `'bgzip -c'` and `'tabix -C -p vcf'`
  - VCF file is converted to FASTA using a combination of `'samtools faidx'` and

    `'bcftools consensus --missing ? -s'`

All different inputs are now in the same format and will follow the same workflow:

1. Custom R script that uses Exon / Intron Locations to splice DNA FASTA (Link to Script )
2. Blast Reference Protein onto spliced DNA FASTA with `'makeblastdb -dbtype nucl'` and

   `'tblastn -seg no -ungapped -comp_based_stats F -outfmt 5'`
3. Custom Python3 script to extract blasted / translated protein and output it in FASTA format (Link to Script).
4. Shell commands to merge together individual proteins into larger datasets (per Protein / per Individual

### Module 3

#### Description of Module:

**Module 3** is designed to perform a phylogenetic analysis, with some modifications specifically designed for palaeoproteomic data. The input of the Module is a FASTA file, containing all of the protein sequences from both the reference dataset and the ancient sample(s) to be analyzed. Accompanying this FASTA file should be a *.TXT file that contains the name of the dataset-FASTA file as well as the names of all of the ancient samples included in that dataset. The dataset will automatically be split into protein specific sub-datasets, each of which will be aligned and checked for SAPs.

The alignment is a two step process which includes first isolating and aligning the modern/reference dataset and then aligning the ancient samples onto the modern ones using Mafft [14]. Isobaric amino acids that cannot be distinguished from each other by the Mass Spectrometer are corrected to ensure the downstream phylogenetic analysis can proceed without problems. Specifically, any time an Isoleucine (I) or a Leucine (L) is identified in the alignment, all of the modern sequences are checked for that position. If all of them share one of the 2 amino acids, then the ancient samples are also switched to that amino acid. If both I and L appear on some present-day samples, both present-day and ancient samples are switched to an L. The user also has the option to provide an additional *.TXT file named 'MASKED'. Using this optional file, the user can 'mask' a present-day sample such that it has the same missing sub-sequences as an ancient sample.

Finally a small report is generated for each ancient sample in the dataset, and a maximum likelihood phylogenetic tree is generated for each protein sub-dataset through PhyML [15]. All protein sub-dataset alignments are then also merged together into a concatenated dataset. The concatenated dataset is used to generate a maximum-likelihood species tree [16] through PhyML and a Bayesian species tree [17, 18] through MrBayes [19] or RevBayes [20]. The tree generation is parallelized using Mpirun [21].

#### Input of the module:

The main input of this module is a FASTA file containing both the ancient sequences to be analysed as well as the full reference data set.

- All proteins for all individuals, the format of fasta sequence labels should be: `>SampleName_ProteinName`
- User must also provide the names of the ancient samples in the analysis. Proteins that are not found in any of the ancient samples but exist inside the input fasta file, will not be included in the analysis.
- Optional - Masking: If the user want to mask a modern sample with the missingness of an ancient one they need to provide a file named 'MASKED' in the module's main directory. This file should contain two columns and as many rows as necessary. Each row should contain the names of two samples, first the name of a modern sample to be masked and second, separated by a whitespace, the name of an ancient sample. The missing positions of each ancient sample will be masked on top of the same positions of the modern sample, 'masking' it as an ancient sample.

#### Workflow:

1. The initial FASTA is split into protein-specific FASTAs with a custom R script (Link to Script).
2. Each protein-specific dataset is aligned using Mafft:
3. Modern and Ancient samples are first separated.

4. Modern samples are aligned with

   ‘mafft --ep 0 --op 0.5 --lop -0.5 --genafpair --maxiterate 20000 --bl 80 --fmodel‘

5. Ancient samples are merged and aligned with modern ones with the ‘mafft-einsi –addlong’ option.
6. Aligned dataset is trimmed of completely missing positions using ‘trimal -noallgaps’
7. A custom R script generates a small table of statistics for each ancient sample in the dataset after correcting for I/L positions (Link to Script).
8. A custom R script concatenates protein-specific FASTAS into Concatenated FASTA. User has the option to filter proteins under certain coverage. A ‘Partition_Helper’ file is also generated, which contains start and stop positions of each protein and can be utilised by NEXUS format phylogenetic software (Link to Script).
9. Masking: Optional step of ’masking’ a modern sample with the missingness of an ancient sample, takes place at this step. This ste will only run if there is file named ’MASKED’ inside the

   /Data_Analysis/ folder .
10. A custom R script is used to convert FASTA files to PHYLIP format (Link to Script).
11. Multithreaded version of PhyML is run on each separate protein-specific data set using:

```
mpirun -n threads phyml-mpi -i dataset.phy -d aa -b 100 -m JTT -a e -s BEST -v e -o tlr
-f m --rand_start --n_rand_starts 4 --r_seed (random_seed_number) --print_site_lnl --print_trac
--no_memory_check
```

12. Concatenated data set is converted to NEXUS format using ‘seqmagick convert –output-format nexus –alphabet protein’ , with minor bash command line fixes to ensure proper formatting.
13. Multithreaded RevBayes is run on the concatenated NEXUS file using a custom RevBayes script Link to Script:
14. Multithreaded version of MrBayes is run on the concatenated NEXUS file using the MrBayes commands:

```
prset aamodelpr = mixed;
mcmc nchains = (number of cores)/4 nruns=(number of cores)/8 ngen = 10000000 samplefreq=100
printfreq=100 diagnfreq=1000;
sumt relburnin = yes burninfrac = 0.25;
sump;
```

   and adding the contenet of the generated ‘Partition_Helper’ file and finally running the file by executing "mpirun -np (number of cores/4) mb-mpi\

### Requirements for running the pipeline

The only true requirement for running any of the 3 modules, is having a Linux machine with Conda installed. All of the required software and packages are downloaded and installed through conda using the provided conda environments. Bellow you can find the full list of software and package used by the pipeline.

#### OS Requirements:

- Linux

#### List of Software and version used by the pipeline:

- Snakemake v7.3.6 [22]
- Conda v22.9.0 [23]
- Samtools v1.15 [10]
- BCFtools v1.15 [24]
- Blast v2.12.0+ [25]
- Angsd v0.937 [26]
- Mafft v7.490 [14, 27]
- Trimal v1.4.rev15 [28]
- Mpirun v4.1.1 [21]
- PhyML v3.3.20200621 [15]
- MrBayes v3.2.7 [19]
- RevBayes v1.0.12 [20]
- Seqmagick v0.8.4 https://github.com/fhcrc/seqmagick
- R v4.1.0 [29]
- Python 3 v3.9.6 [30]

#### R packages

- Bioconductor - ShortRead v1.50.0 [31]
- Phyclust v0.1.30 [32]
- Stringr v1.4.0 [33]

#### Python packages

- Biopython v1.79 [34]
- OS package [30]
- Sys package [30]
- Requests package v2.26.0 [35]
- RE package v2.2.1[36]

### Palaeo proteomic hominid reference dataset

This section of the supplementary file is dedicated to describing how the 'Palaeo proteomic hominid reference dataset' was generated. The dataset is available here

#### Choosing and preparing the list of proteins.

We selected 6 publications cataloging proteins identified in either teeth or bone tissue[37, 38, 39, 40, 41, 42]. From these publications we compiled a list of 1696 unique protein names, which are provided in this file in the main Github repository and can also be found within the Zenodo database: as `\Reference_Protein_List.txt"` This file is a tab separated list of every protein generated for the 'Reference Dataset'. Each protein is linked to at least one publication where it was identified in either bone or tooth tissue. If multiple publications mention a protein, the names of these publications are separated with a comma (' , '). We modified this list and used it as an input for Module 1 of the pipeline. From these 1696 proteins, 153 could not be matched to an Ensembl Gene ID. This left a total of 1543 proteins that were successfully translated.

#### Choosing and preparing samples for translation.

Samples were chosen and used from the following publications:

– (1) 1KG high coverage[43].
– (2) Great ape genomes project[44].
– (3) Morphometric, Behavioral, and Genomic Evidence for a New Orangutan Species[45].
– (4) A high-coverage Neandertal genome from Vindija Cave in Croatia[46].
– (5) A high-coverage Neandertal genome from Chagyrskaya Cave[47].

#### Choosing individuals for the data set:

For publications 2,3 and 5 all available samples were used. For publication 1, only a maximum of 3 individuals from each labeled population were used. For publication 4 , the individual named 'Mezmaiskaya' was removed from the data set due to a high amount of predicted unique SAPs. We believe that due to the low coverage of the individual, a high number of variants might be miss-called. A table with the full table/list of individuals included in the dataset can be found in the main page of the Github here. The table connects the name of each sample with the species or population it belongs to, the publication the data originates from and the type of data that was used for the translation of that sample (e.g. VCF or BAM files).

#### Reference Genomes:

We chose to use the human reference genome as the basis for our translations, due to its higher level of annotation. For this purpose, all individuals were mapped onto either GRCh37[48] or GRCh38[49]. Individuals from datasets 1,4 and 5 were already mapped onto a human reference genome. For datasets 2 and 3, raw fastq files were downloaded and mapped onto the GRCh38 human reference. The mapping workflow is provided in the form of a snakemake python script along with a Conda environment that contains all the necessary software to run it. The re-mapped bam files are available upon request.

##### Final execution:

Once the BAM and VCF files were prepared, they were utilized as input for Module 2 and the targeted proteins were translated for every chosen individual. The process of translating these datasets is described in detail in the github tutorial, in “STEP 2”, including the preprocessing that some of the VCF files need to go through. We used GRCh38 as the annotation reference for datasets 1,2 and 3 and GRCh37 for datasets 4 and 5. The generated protein fasta files were collected from the output folder and assembled into the final folder, which is available in Zenodo.

references

[32] Wei-Chen Chen. “Overlapping codon model, phylogenetic clustering, and alternative partial expectation conditional maximization algorithm”. PhD thesis. Iowa State University, 2011.

[33] Hadley Wickham. *stringr: Simple, Consistent Wrappers for Common String Operations*. http://stringr.tidyverse.org https://github.com/tidyverse/stringr. 2022.

[34] Peter JA Cock et al. “Biopython: freely available Python tools for computational molecular biology and bioinformatics”. In: *Bioinformatics* 25.11 (2009), pp. 1422–1423.

[35] Rakesh Vidya Chandra and Bala Subrahmanyam Varanasi. *Python requests essentials*. Packt Publishing Ltd, 2015.

[36] Guido Van Rossum. *The Python Library Reference, release 3.8.2*. Python Software Foundation, 2020.
